# USP1 inhibition promotes RAD18-dependent PCNA degradation and BRCA1 synthetic lethality

**DOI:** 10.64898/2026.02.25.708068

**Authors:** Nicholas W. Ashton, Ramya Ravindranathan, Emilie J. Korchak, Ozge S. Somuncu, Gabriella A. Zambrano, Sirisha V. Mukkavalli, Shuhei Asada, Dmitry Korzhnev, Irina Bezsonova, Alan D. D’Andrea

## Abstract

The proliferating cell nuclear antigen (PCNA) sliding clamp is mono-ubiquitinated by RAD6-RAD18 in response to DNA damage, initiating the DNA damage tolerance pathway of translesion synthesis. The molecular basis by which RAD18 engages PCNA has, however, remained incompletely defined. Mono-ubiquitinated PCNA is subsequently poly-ubiquitinated with K48-linked chains that target PCNA for degradation. Ubiquitin-specific protease 1 (USP1) reverses PCNA mono- and poly-ubiquitination; accordingly, inhibiting USP1 causes the accumulation of mono-ubiquitinated PCNA at replication forks and a reduction in total PCNA levels. USP1 inhibitors promote the accumulation of ssDNA gaps (ssGAPs) in newly replicated DNA and are synthetic lethality in BRCA1-deficient cells. Here, we combine computational and structural approaches to identify and characterize a PCNA-interacting peptide (PIP) motif in RAD18. This PIP motif is required for RAD18-dependent DNA damage-induced PCNA ubiquitination and PCNA turnover. Mutation of the RAD18-PCNA interface reduces ssGAP accumulation and USP1 inhibitor sensitivity in BRCA1-deficient cells. Furthermore, cells adapted to prolonged USP1 inhibition exhibit reduced RAD18 levels, suggesting that deregulation of RAD18 contributes to a biologically relevant drug resistance mechanism. This resistance could be overcome by inhibiting the Ataxia telangiectasia and Rad3-related (ATR) kinase. Together, these findings define a molecular interface required for RAD18-dependent PCNA mono-ubiquitination and identify it as a key determinant of USP1-BRCA1 synthetic lethality.

## Introduction

DNA replication and repair require tightly coordinated DNA synthesis. The homotrimeric proliferating cell nuclear antigen (PCNA) sliding clamp is a key component of these pathways in eukaryotes. PCNA encircles double-stranded DNA and stabilizes most DNA polymerases on the template strand (1). This includes the replicative polymerases Pol δ (2, 3) and Pol ε (4). PCNA also functions as an organizing hub for other replication-related proteins. For instance, during discontinuous lagging strand synthesis, PCNA acts as a platform for Pol δ, flap endonuclease 1 (FEN1) (5) and DNA ligase 1 (LIG1) (6, 7), to coordinate the maturation of Okazaki fragments (8).

Most proteins bind PCNA via eight-amino acid PCNA-interacting protein (PIP) motifs that interact with the universal binding site on the front surface of each PCNA monomer (9). While the precise sequences of these motifs vary considerably, most are recognizable variants of a common consensus sequence (QxxΦxxΨΨ; Φ = hydrophobic, Ψ = aromatic). Due to the large number of proteins that must compete for binding to the same site on PCNA, cells require finely tuned and multilayered regulatory mechanisms to spatially and temporally control the access of different partners (10). The DNA damage-induced recruitment of the Y-family translesion polymerases Pol ι, Pol η, Pol κ and Rev1 is a notable example, driven by mono-ubiquitination of PCNA at lysine residue 164 (K164). Mono-ubiquitination creates an additional binding platform that these polymerases interact with through their ubiquitin-binding domains (11–13). TLS polymerases feature comparatively open active sites, allowing them to bypass a range of DNA damage adducts, including UV-induced photoproducts (14). Recent studies suggest that RAD18-dependent PCNA mono-ubiquitination also contributes to post-replicative TLS at ssDNA gaps, known to arise during replication in BRCA1/2-deficient cancer cells (15, 16).

Protein ubiquitination occurs through a multi-step enzymatic pathway, in which a ubiquitin molecule is initially bound by an E1 ubiquitin activating enzyme, before being transferred to an E2 conjugating enzyme. The E2, in complex with an E3 ligase, then catalyzes the formation of an isopeptide bond between ubiquitin and the target lysine residue of the substrate protein (17). Members of the RING finger E3 ligase family do not form catalytic intermediates with ubiquitin, but rather provide substrate specificity to the complex by guiding the ubiquitin-laden E2 to the target lysine residue of the substrate protein (18). The RAD6(E2)-RAD18(E3) complex is the predominant E2-E3 complex responsible for the mono-ubiquitination of PCNA at lysine residue 164 (K164) in response to DNA damage (19). Biochemical studies have demonstrated that RAD18 molecules dimerize via their RING domains, and that RAD6 interacts with one of these monomers to form a functional RAD6-RAD18_2_ complex (20–22). RAD18 is recruited to single-stranded DNA at stalled replication forks via a direct interaction with the replication protein A subunit, RPA70 (23–25). Precisely how RAD18 interacts with PCNA to direct RAD6-mediated ubiquitination has however remained unclear. Indeed, while a C-terminal truncation mutant of RAD18 (residues 16-366) is known to form a complex with RAD6 and PCNA, this interaction has not been localized to a specific region of RAD18 (22). It has however been proposed that RAD18 likely interacts with the universal-binding site of PCNA, as the ability of RAD6-RAD18_2_ to mono-ubiquitinate PCNA *in vitro* is suppressed by the presence of Pol η or Pol δ, suggesting that RAD18 competes with these polymerases for binding to the same site on PCNA (26).

The deubiquitinating enzyme, Ubiquitin Specific Protease 1 (USP1) and its cofactor, USP1-associated factor 1 (UAF1) (27), catalyze the deubiquitination of mono- and poly-ubiquitinated PCNA (28, 29). USP1 is required for fork protection, and its genetic loss is synthetically lethal with BRCA1 deficiency (30). Numerous allosteric USP1 inhibitors have been developed for the treatment of BRCA1-deficient cancers (31–35). USP1 inhibitors cause an accumulation of mono-ubiquitinated PCNA at replication forks, as well as a concurrent reduction in total PCNA levels (31, 32). Cell killing by USP1 inhibitors correlates with the generation of single-strand DNA gaps (ssGAPs) in cells (32). Importantly, the accumulation of these ssGAPs, as well as the synthetic lethality between USP1 inhibition and BRCA1 deficiency, can be reduced by silencing of RAD18 (30–32, 36).

In this work, we identify a non-consensus PIP motif within the SAP domain of RAD18 and demonstrate that the interaction of this motif with PCNA is required for DNA damage-induced PCNA mono-ubiquitination. Mutating the PIP motif reduces the synthetic lethality of USP1 inhibitors in BRCA1-deficient cells. Together, these findings define a RAD18-PCNA interface required for PCNA mono-ubiquitination and establish its role in USP1-BRCA1 synthetic lethality.

## Results

### USP1 reduces RAD18-mediated PCNA mono-ubiquitination and PCNA degradation

Previous works have established that PCNA is mono- and poly-ubiquitinated at lysine residue 164 via a two-step enzymatic process. PCNA is initially mono-ubiquitinated by RAD6-RAD18_2_, and subsequently poly-ubiquitinated by other E2-E3 pairs (**Fig. 1A**). The USP1-UAF1 complex deubiquitinates mono- and poly-ubiquitinated PCNA (28, 29). USP1 has therefore been suggested to prevent RAD18-dependent proteasomal degradation of poly-ubiquitinated PCNA (31, 36).

**Figure 1.**
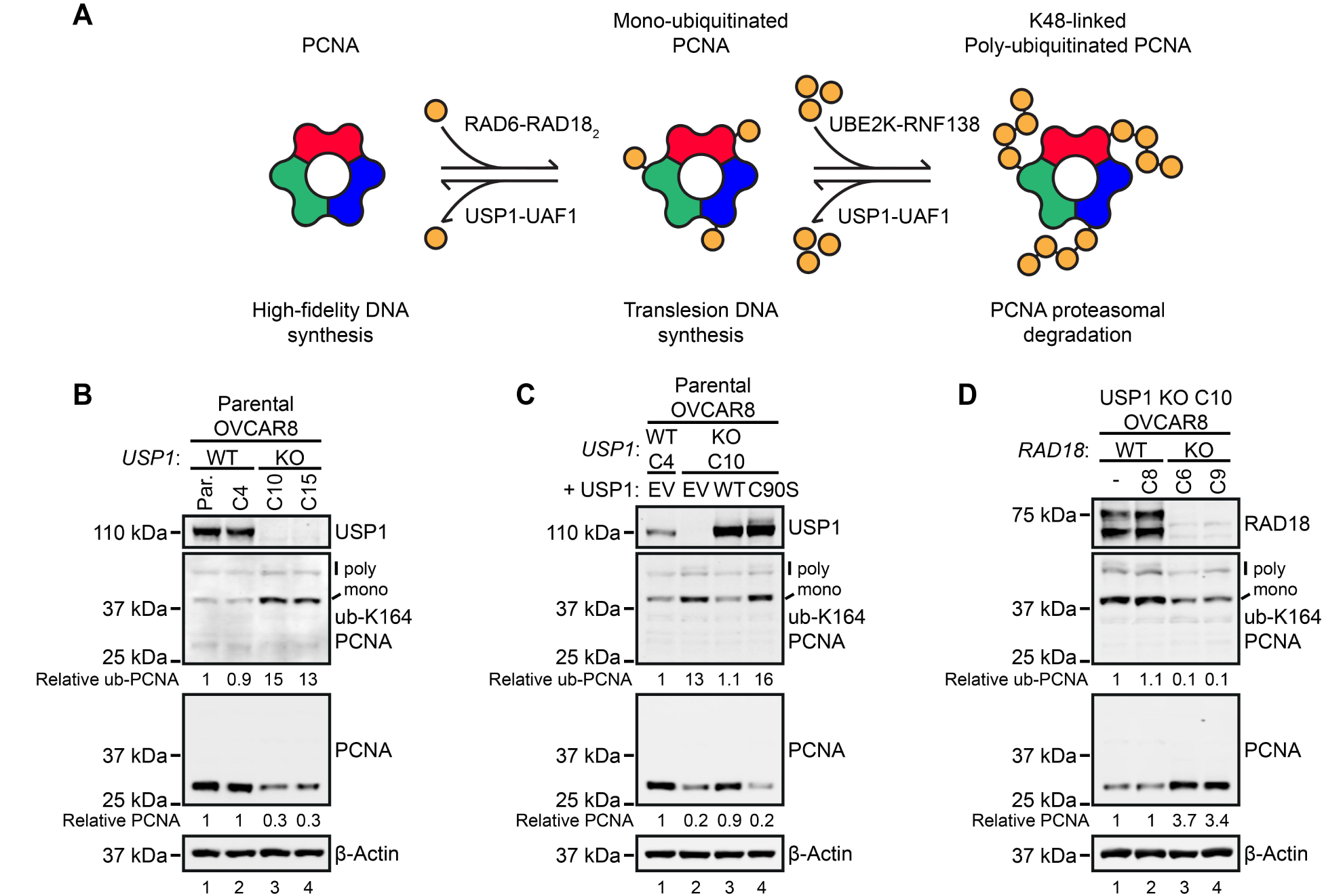
USP1 reduces RAD18-mediated PCNA mono-ubiquitination and PCNA degradation. (**A**) Schematic illustrating PCNA ubiquitination and deubiquitination pathways. (**B**) Immunoblot analysis of whole-cell lysates from parental OVCAR8 cells (Par.), a WT USP1 clone (C4), and two USP1 knockout clones (C10 and C15). Relative total PCNA levels were determined from the PCNA:β-actin ratio, and relative mono-ubiquitinated PCNA (ub-PCNA) levels were determined from the ub-PCNA:PCNA ratio. Values are expressed relative to parental OVCAR8 cells (lane 1). (**C**) Immunoblot analysis of ubiquitinated and total PCNA in whole-cell lysates from a WT USP1 OVCAR8 clone (C4) and a USP1 KO clone (C10) stably expressing WT USP1, catalytically inactive USP1 (C90S), or empty vector (EV). Relative PCNA and ub-PCNA levels were calculated as described in (B). (**D**) Immunoblot analysis of whole-cell lysates from WT RAD18 (C8) or RAD18 knockout clones (C6 and C9) generated in a USP1 KO OVCAR8 background (C10). Relative PCNA and ub-PCNA levels were calculated as described in (B).

To study the role of RAD18 in this context, we used CRISPR-Cas9 to target USP1 in the ovarian cancer cell line, OVCAR8. Immunoblotting of knockout (KO) clones revealed an increase in mono- and poly-ubiquitinated PCNA, as well as a ≈ 70% reduction in total PCNA levels (**Fig. 1B**), consistent with the model shown in **Fig. 1A**. To ensure these effects were due to loss of USP1 enzymatic activity, we used lentiviral transduction to stably express WT or catalytically inactive USP1 (C90S) in a USP1 KO clone. Re-expressing WT, but not C90S USP1, restored PCNA to levels observed in WT USP1 cells (**Fig. 1C**). We also used CRISPR Cas9 to generate RAD18 KO clones in a USP1 KO background. Strikingly, loss of RAD18 led to a 3-4-fold increase in PCNA levels, and a reduction in PCNA mono- and poly-ubiquitination (**Fig. 1D**). In contrast, KO of RAD18 in parental OVCAR8 cells led to a reduction in PCNA mono-ubiquitination, while levels of total and poly-ubiquitinated PCNA were unaltered (**Fig. S1**). These data support a model in which RAD18 promotes PCNA turnover, however, this activity is normally suppressed by USP1.

### The RAD18 SAP domain interacts with PCNA

RAD18 appears to drive PCNA turnover by mediating PCNA mono-ubiquitination and creating a substrate that can be poly-ubiquitinated by other E2-E3 pairs. To test this hypothesis, we sought to create a separation-of-function point mutant of RAD18 that cannot direct PCNA mono-ubiquitination. The SAP domain of RAD18 is essential for the RAD6-RAD18_2_-mediated mono-ubiquitination of PCNA *in vitro* and in cells (**Fig. 2A**) (21, 22, 37). As with SAP domains from other proteins, the RAD18 SAP domain possesses residual DNA-binding activity (22, 37). DNA-binding is not, however, required for RAD18 function, as SAP domain deletion mutants still accumulate at sites of DNA damage (38, 39). Furthermore, the SAP domain of RAD18 is required for PCNA mono-ubiquitination *in vitro* even in the absence of DNA (37). Previous works have therefore hypothesized that the RAD18 SAP domain contains an unidentified PCNA-binding interface (37, 40).

**Figure 2.**
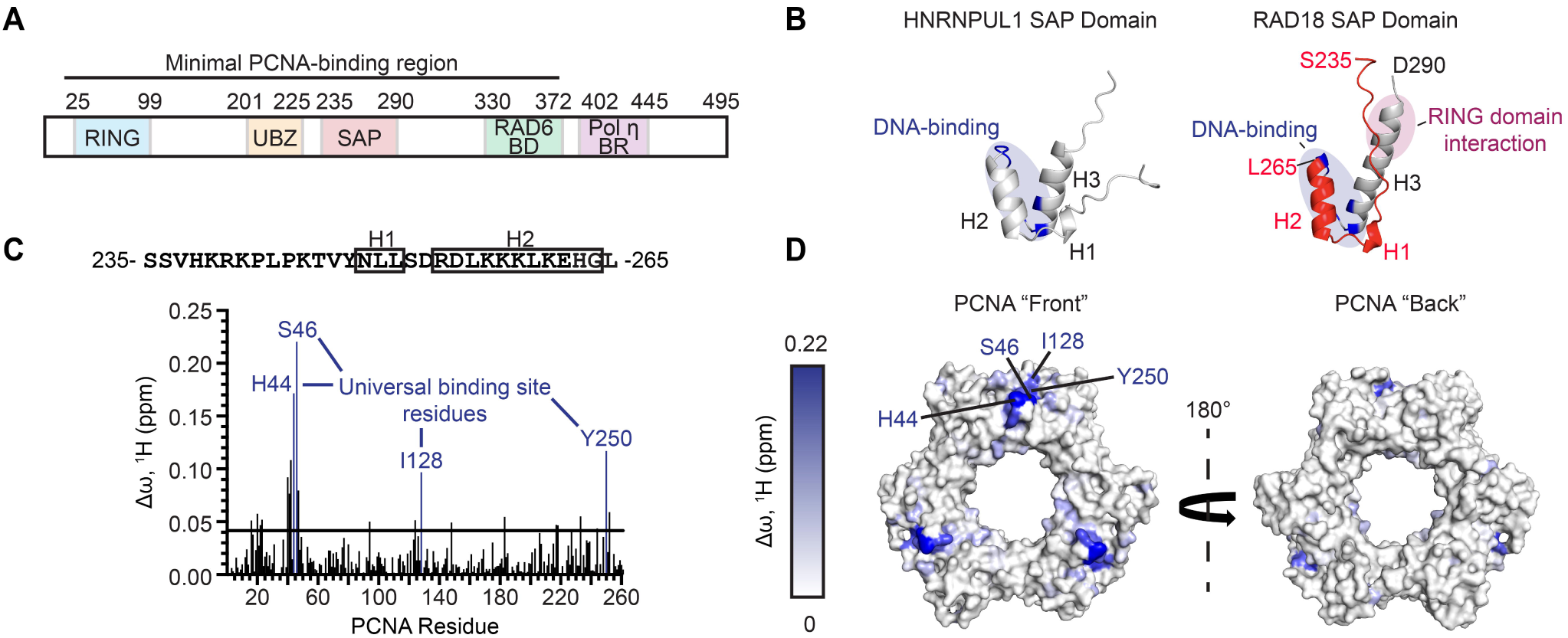
The RAD18 SAP domain interacts with PCNA. (**A**) Schematic of RAD18 domain architecture, including the RING domain, ubiquitin-binding zinc finger (UBZ), SAP domain, RAD6-binding domain (RAD6 BD), and Pol η-binding region (Pol η BR). (**B**) Solution structure of the HNRNPUL1 SAP domain (PDB: 1ZRJ; left) and an AlphaFold model of the RAD18 SAP domain (right). Helices 1–3 are labeled (H1-H3). (**C**) Bar graph showing NMR chemical shift perturbations (Δω) in the ^15^N-TROSY spectrum of ^15^N/^2^H-labeled PCNA following addition of unlabeled WT RAD18 peptide (residues 235-265). Residues comprising the PCNA universal binding site are highlighted in blue. (**D**) Mapping of the RAD18 peptide (235–265) binding interface onto the surface of PCNA (PDB: 4RIF), color-coded according to Δω values from (C), from lowest (white) to highest (blue). Universal binding site residues are labeled. Structures are shown in two orientations (180° rotation)

SAP domains are characterized by a stable helix (H2)-loop-helix (H3) core, with an additional smaller N-terminal helix (H1) located within the unstructured N-terminal portion of the domain (41). To visualize the RAD18 SAP domain, we accessed an AlphaFold model of RAD18 from the AlphaFold Protein Structure Database (42) and isolated residues S235-D290 (**Fig. S2**). The predicted SAP domain structure resembles the human heterogeneous nuclear ribonucleoprotein U-like protein SAP domain, (PDB: IZRJ) with the conserved DNA-binding residues located on analogous surfaces of helices H2 and H3, and on the connecting loop (22, 37) (**Fig. 2B**). The full-length RAD18 model also predicts an interaction between H3 of the SAP domain and a helix within the RAD18 RING domain, consistent with recent experimental observations (**Fig. S2**) (40).

To test whether the RAD18 SAP domain binds PCNA, we used solution nuclear magnetic resonance (NMR) spectroscopy. A peptide spanning RAD18 residues S235-L265 – encompassing the flexible N-terminal segment, helices H1 and H2, and the intervening loop – was titrated into ^15^N/^2^H-labeled PCNA, and the resulting residue-specific chemical shift perturbations (Δω) in the ^15^N TROSY spectrum were quantified (**Fig. 2C**). As the NMR resonances of PCNA were previously assigned (BMRB 15501) (43), the Δω values could be mapped onto the PCNA structure (PDB: 4RJF) (44) to define the peptide-binding surface (**Fig. 2D**). The largest chemical shift perturbations were observed for residues H44, S46, I128, and Y250, corresponding to the well-described universal-binding site of PCNA (9, 45). These data indicate that the RAD18 SAP domain peptide interacts with the PCNA universal-binding site.

### A non-consensus PIP motif mediates RAD18-dependent PCNA mono-ubiquitination

PCNA-binding proteins typically interact with the universal-binding site of PCNA via 8-amino acid PIP motifs that form a short 3_10_ helix (9). Although RAD18 lacks a consensus PIP motif sequence within the RAD18 SAP domain, or elsewhere within the protein, we identified a well-conserved sequence resembling the non-canonical PIP motifs of translesion polymerases (e.g. Pol η, Pol ι) (**Fig. 3A**). Such motifs often lack a glutamine in position 1 and contain bulky non-aromatic hydrophobic residues in positions 7-8 (46). Notably, residues 4-7 of the putative motif correspond to the small H1 helix of the SAP domain, consistent with the structure of known PIP motifs (9). Consistent with this interpretation, AlphaFold modelling of residues K241-R254 in complex with PCNA suggested a binding mode compatible with engagement of the PCNA universal-binding site and resembling known PIP-PCNA interactions (**Fig. 3B**).

**Figure 3.**
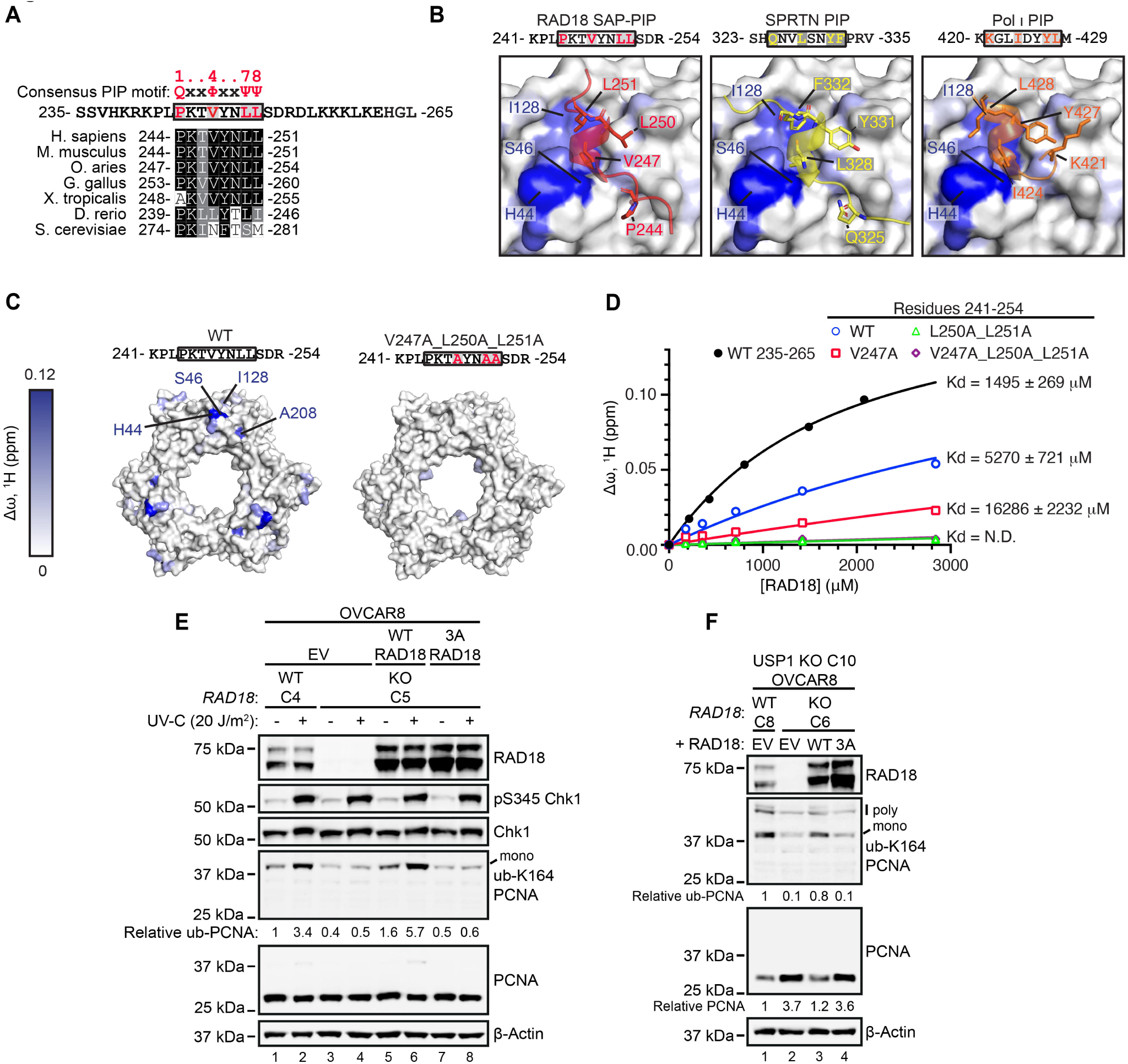
A non-consensus PIP motif mediates RAD18-dependent PCNA mono-ubiquitination. (**A**) Sequence alignment of a putative PIP motif within SAP domains of RAD18 homologs. Conserved residues are shaded (black = identical; gray = similar; white = dissimilar). Q = glutamine; Φ = hydrophobic; Ψ = aromatic. (**B**) AlphaFold3 model of the RAD18 SAP domain PIP motif bound to the PCNA universal binding site (left). Structures of SPRTN (middle) and Pol ι (right) PIP motifs bound to PCNA were derived from crystal structures (PDB: 5IY4 and 2ZVM, respectively). Side chains of PIP motif residues 1, 4, 7, and 8 are shown. The colored PCNA structure from (Fig. 2D) was superimposed onto each model. (**C**) Mapping of PCNA-binding surfaces for WT and V247A_L250A_L251A RAD18 peptides (residues 241-254) using NMR Δω values, displayed on the PCNA surface (PDB: 4RIF). Universal binding site residues are highlighted. (**D**) NMR titration analysis comparing PCNA binding to the longer (residues 235-265) and shorter RAD18 peptides (residues 241-254; WT, V247A, L250A_L251A and V247A_L250A_L251A). Binding affinities were assessed by monitoring a representative PCNA peak during peptide titration. (**E**) Immunoblot analysis of UV-C-induced PCNA mono-ubiquitination in WT RAD18 OVCAR8 cells (C4) and RAD18 KO OVCAR8 cells (C5) stably expressing WT RAD18, 3A RAD18 (V247A_L250A_L251A), or empty vector (EV). Cells were exposed to 20 J m⁻² UV-C and harvested after 2 h. Relative mono-ubiquitinated PCNA (ub-PCNA) was calculated from the ub-PCNA:PCNA ratio and normalized to untreated WT RAD18 cells (lane 1). (**F**) Immunoblot analysis of whole-cell lysates from WT RAD18 (C8) and RAD18 KO (C6) clones generated in a USP1 KO OVCAR8 background (C10). RAD18 KO cells were complemented with WT RAD18, 3A RAD18, or EV. Relative total PCNA levels were determined from PCNA:β-actin ratios, and ub-PCNA levels were calculated from ub-PCNA:PCNA ratios.

To verify whether this putative PIP motif interacts with PCNA, we repeated the NMR titration experiments using a shorter RAD18 peptide spanning residues K241-R254, encompassing this motif (P244-L251) and its flanking residues. In addition to a WT peptide, we tested variants in which residues 4, 7 and 8 of the PIP motif were substituted with alanine (V247A, L250A, L251A). The resulting chemical shift perturbations in the PCNA spectra were quantified and mapped onto the PCNA structure (**Fig. 3C and Fig. S3**). These analyses revealed binding between the shorter WT peptide and the universal-binding site of PCNA, which was disrupted by the V247A, L250A and L251A mutations. Although the shorter WT peptide bound to PCNA less readily than the longer peptide (S235-L265) (**Fig. 3D**), this is consistent with other PIP motif-PCNA interactions, in which flanking residues can stabilize the PIP helix or contribute additional contacts with PCNA (9).

To test the requirement of the RAD18 PIP motif for PCNA mono-ubiquitination, we stably expressed WT RAD18 or the 3A RAD18 mutant, in RAD18 KO OVCAR8 cells. Following UV-C irradiation, RAD18 KO cells failed to induce PCNA mono-ubiquitination, and this defect was rescued by WT but not 3A RAD18 (**Fig. 3E**). We observed the same requirement for the SAP domain PIP motif in RAD18 KO RPE-1 cells (**Fig. S4**). To study the role of RAD18-mediated mono-ubiquitination in PCNA turnover, we also expressed WT or 3A RAD18 in RAD18-USP1 double KO OVCAR8 cells. Expression of WT RAD18 strongly reduced total PCNA levels and increased mono-ubiquitinated PCNA, resembling USP1 KO cells, while expression of 3A RAD18 had no effect on either ubiquitinated or total PCNA (**Fig. 3F**). Together, these data establish that the SAP domain PIP motif forms a PCNA-binding interface required for PCNA mono-ubiquitination and subsequent PCNA turnover in the absence of USP1.

A recent study reported an additional putative PIP motif within the Pol η-binding region of the unstructured RAD18 C-terminus (residues Q404-E411) (47), adjacent to ATR- and JNK-phosphorylation sites (47, 48) (**Fig. S5A-B**). Similar to the SAP domain motif, this C-terminal sequence poorly conforms to the consensus PIP sequence, not only lacking bulky aromatic residues in positions 7 and 8, but also containing a charged glutamic acid in position 8. Although this region is outside the minimal PCNA-binding region of RAD18 (residues 16-366) (22), mutating Q404 and L407 (positions 1 and 4) in this motif to alanine was found to impair RAD18-mediated PCNA mono-ubiquitination *in vitro*. To assess the relevance of this region in our system, we introduced a Q404A_L407A RAD18 mutant into RAD18 KO OVCAR8 cells, and measured UV-C-induced PCNA mono-ubiquitination (**Fig. S5C**). These mutations indeed diminished PCNA mono-ubiquitination by ≈ 50% relative to the over-expressed WT protein, consistent with a role for this region in PCNA mono-ubiquitination. Given that mutation of the SAP domain PIP motif caused a near-complete loss of PCNA mono-ubiquitination, however, we used the SAP 3A mutant in subsequent experiments to selectively and robustly disrupt RAD18-directed PCNA mono-ubiquitination.

### RAD18-dependent PCNA mono-ubiquitination and degradation enables USP1 inhibitor-mediated synthetic lethality in BRCA1-deficient cells

The genetic depletion or inhibition of USP1 in BRCA1-deficient cells results in the accumulation of replication defects and cell death, in a RAD18-dependent manner (30, 32). To study the role of PCNA-binding by RAD18 in this context, we firstly used CRISPR-Cas9 to generate RAD18 KO clones of the BRCA1-null ovarian cancer cell line, UWB1.289 (**Fig. 4A**). This cell line carries a germline heterozygous frameshift mutation in BRCA1 and loss of heterozygosity of the WT allele (49). Consistent with our previous studies with RAD18 siRNA (30, 32), RAD18 KO conferred resistance of these cells to two allosteric USP1 inhibitors, I-138 and KSQ-4279 (**Fig. 4B-C and Fig. S6**). We stably reconstituted a RAD18 KO clone (C13) with wild-type RAD18 or the RAD18 3A mutant using lentiviral transduction (**Fig. 4D**).

**Figure 4.**
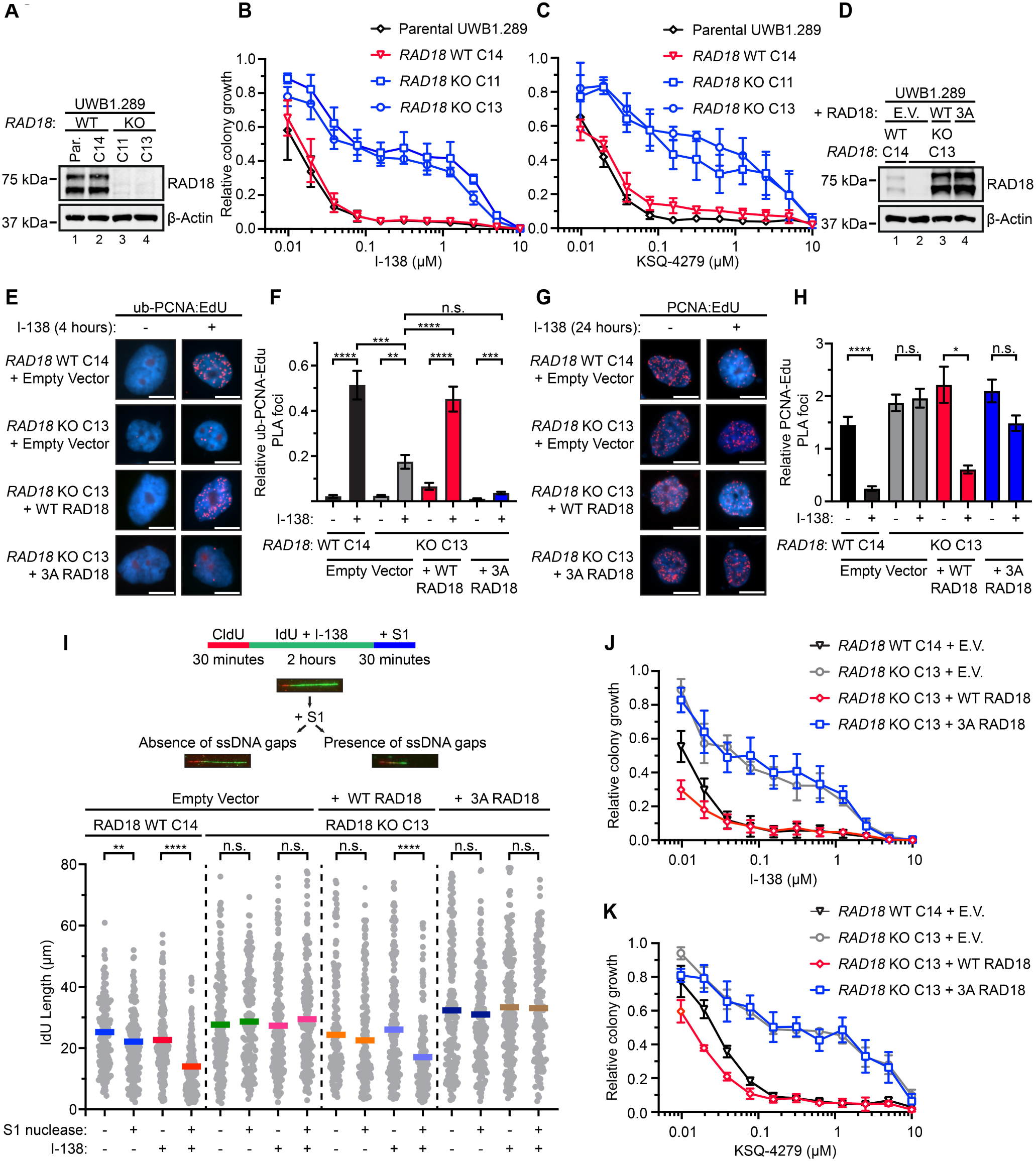
RAD18-dependent PCNA mono-ubiquitination and degradation enables USP1 inhibitor-mediated synthetic lethality in BRCA1-deficient cells. (**A**) Immunoblot analysis of whole-cell lysates from parental UWB1.289 cells (Par.), a WT RAD18 clone (C14), and two RAD18 knockout clones (C11 and C13). (**B-C**) Clonogenic survival of the cell lines in (A) following treatment with the USP1 inhibitors I-138 (B) or KSQ-4279 (C). WT RAD18 cells were stained after 10 days of treatment, while RAD18 KO clones were stained after 14 days. Values represent means of three biological replicates ± SD. (**D**) Immunoblot analysis of whole-cell lysates from a WT RAD18 UWB1.289 clone (C14) and a RAD18 KO clone (C13) stably expressing WT RAD18, 3A RAD18 (V247A_L250A_L251A), or empty vector (EV). (**E**) Representative SIRF images detecting ub-PCNA:EdU PLA foci at replication forks following treatment with DMSO or I-138 for 4 h. Scale bar, 5 μm. (**F**) Quantification of ub-PCNA:EdU PLA foci. Error bars represent SEM. Statistical analysis was performed using a Mann-Whitney test (n.s. = not significant; *P < 0.05; **P < 0.01; ***P < 0.001; ****P < 0.0001). (**G**) Representative SIRF images detecting PCNA:EdU PLA foci following treatment with DMSO or I-138 for 24 h. Scale bar, 5 μm. (**H**) Quantification of PCNA:EdU PLA foci. Statistical analysis as in (F). (**I**) Top, schematic of DNA fiber assay with or without S1 nuclease treatment. Bottom, quantification of IdU tract lengths from the cell lines in (D) following 2 h treatment with 0.5 μM I-138 ± S1 nuclease. Each data point represents an individual fiber; horizontal lines indicate median tract length. Statistical analysis was performed using a Mann-Whitney test (n.s. = not significant; **P < 0.01; ****P < 0.0001). (**J-K**) Clonogenic survival of the cell lines in (D) following treatment with I-138 (J) or KSQ-4279 (K). Total colony volume was normalized to DMSO-treated controls. Values represent means of three biological replicates ± SD.

RAD18 is recruited by RPA to ssDNA at replication forks, where it directs the mono-ubiquitination of PCNA (23). To specifically measure PCNA mono-ubiquitination and turnover at these sites, we used an imaging-based proximity ligation technique – quantitative in situ analysis of protein interactions at DNA replication forks (SIRF) (50) – to specially detect fork-associated PCNA. As expected, treatment of a WT UWB1.289 clone (C14) with I-138 led to the rapid accumulation of ubiquitinated PCNA at forks within 4 hours, which was not observed in a RAD18 KO clone (C13) (**Fig. 4E-F**). This accumulation was rescued by the stable expression of WT RAD18, although not by the 3A RAD18 mutant. We also used the SIRF assay to measure the loss of fork-associated PCNA at a later time-point. In WT cells, as well as in the RAD18 KO cells reconstituted with WT RAD18, we observed a strong reduction in fork-associated PCNA after 24 hours of I-138 treatment (**Fig. 4G-H**). In contrast, we did not detect a statistically significant reduction in PCNA at replication forks in RAD18 KO cells, or in those cells reconstituted with 3A RAD18.

USP1 inhibitors induce replication defects in sensitive cell lines, as observed by the rapid accumulation of ssDNA gaps (ssGAPs), which are detected by S1 nuclease-treated DNA fiber assays (32). To assess the role of RAD18-mediated PCNA mono-ubiquitination in this process, we quantified ssGAPs in cells expressing WT and 3A RAD18 (**Fig. 4I**). Consistent with our previous findings, I-138 treatment induced an accumulation of ssGAPs in WT UWB1.289 cells, but not in RAD18 KO cells, as indicated by reduced IdU tract length following S1 nuclease treatment. Importantly, the accumulation of these gaps was restored in RAD18 KO cells expressing WT but not 3A RAD18.

Finally, to determine the requirement of the RAD18 PIP-PCNA interface in USP1 inhibitor-induced cell killing, we assessed the sensitivity of these cell lines to the USP1 inhibitors, I-138 and KSQ-4279. While the stable expression of WT RAD18 restored the sensitivity of RAD18 KO UWB1 cells to I-138 and KSQ-4279, cells expressing 3A RAD18 resembled RAD18 KO cells (**Fig. 4J-K**). Together, these data indicate that RAD18-dependent PCNA mono-ubiquitination is a central molecular step linking PCNA turnover, replication defects, and USP1-inhibitor sensitivity in BRCA1-deficient cells.

### USP1 inhibitor-resistant UWB1 cells exhibit attenuated PCNA ubiquitination

Many cancers that initially respond to treatment ultimately develop drug resistance through selection of clones with pre-existing or acquired alterations in drug targets or effector pathways (51). We therefore hypothesized that deregulation of RAD18 and/or PCNA mono-ubiquitination may occur in BRCA1-null cell lines following long-term exposure to USP1 inhibitors. To test this, we cultured UWB1.289 cells in progressively increasing concentrations of I-138 or KSQ-4279 for ≈ 4 months to generate adapted populations (I-138_R and KSQ_R). As controls, we maintained parallel cultures in equivalent volumes of DMSO (DMSO_I and DMSO_K) (**Fig. 5A**). Both adapted populations acquired comparable resistance to I-138 and KSQ-4279, whereas control cells remained highly sensitive (**Fig. 5B and Fig. S7A**).

**Figure 5.**
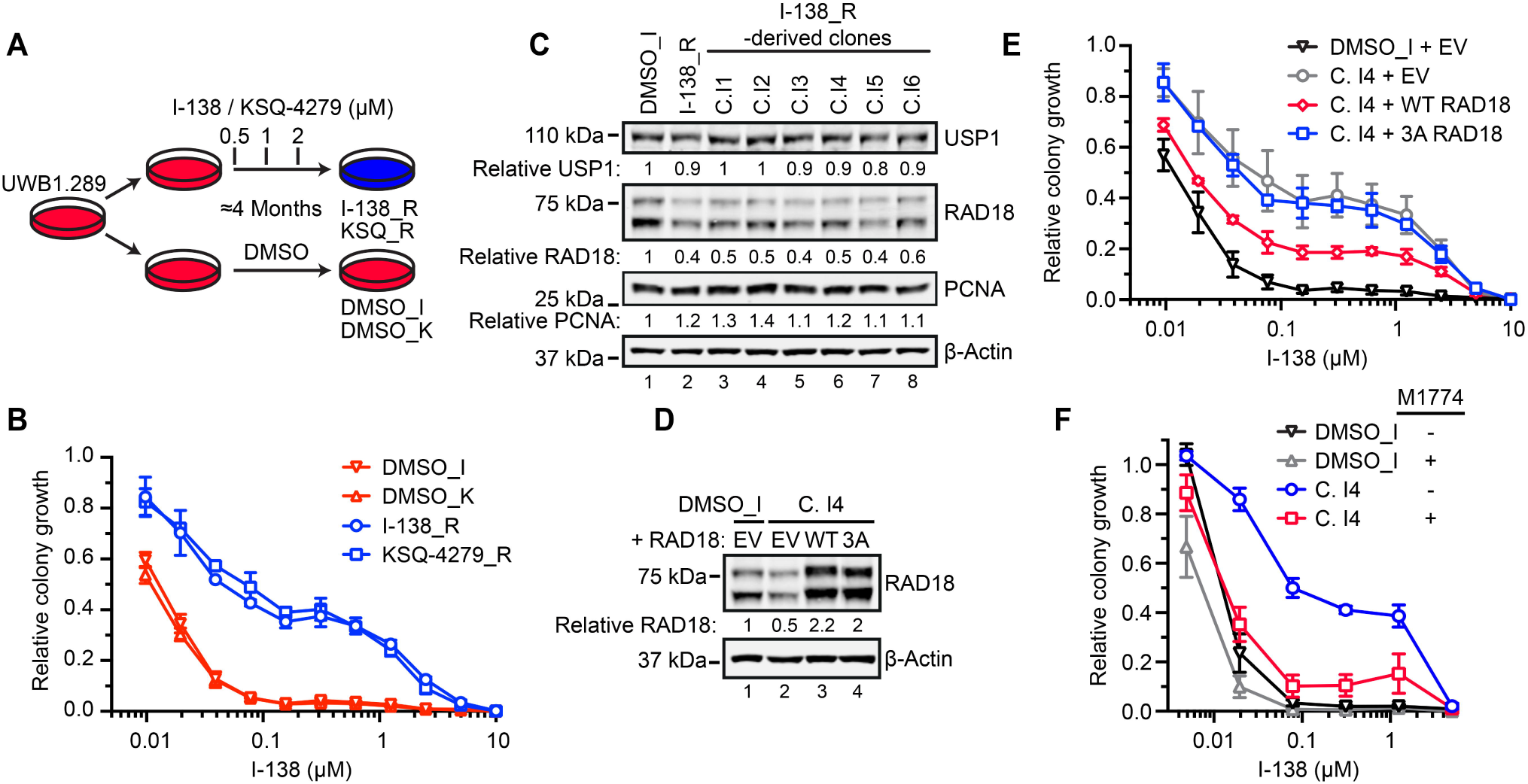
USP1 inhibitor-resistant UWB1 cells exhibit attenuated PCNA ubiquitination. (**A**) Schematic illustrating adaptation of UWB1.289 cells to increasing concentrations of USP1 inhibitors I-138 or KSQ-4279, generating I-138_R and KSQ_R populations, respectively. Parallel cultures treated with DMSO alone are designated DMSO_I and DMSO_K. (**B**) Clonogenic survival of the populations in (A) following treatment with I-138. Adapted populations were maintained drug-free for ≥1 week before analysis. Values represent means of three biological replicates ± SD. (**C**) Immunoblot of whole-cell lysates from DMSO_I, I-138_R, and derived clones I1-I6. USP1, RAD18, and PCNA levels were normalized to β-Actin and expressed relative to lane 1. (**D**) Immunoblot of DMSO_I or C.I4 cells stably expressing WT RAD18, 3A RAD18, or empty vector. (**E**) Clonogenic survival of cell lines in (D) following treatment with I-138. Values represent means of three biological replicates normalized to DMSO controls ± SD. (**F**) Clonogenic survival of DMSO_I and C.I4 cells seeded in media with or without 0.5 nM M1774 and treated with I-138. Values represent means of three biological replicates normalized to DMSO controls ± SD.

We isolated six clonal derivatives from each adapted cell line (I1-I6 and K1-K6), all of which displayed similar resistance to USP1 inhibitors (**Fig. S7B-C**). Immunoblotting revealed reduced RAD18 levels in both adapted polyclonal populations and all individual clones (**Fig. 5C and Fig. S7D**), with some clones also showing modest increases in total PCNA and reduced USP1 levels. Stable expression of WT, but not 3A RAD18, in a representative resistant clone (C.I4), partially restored sensitivity to I-138 and KSQ-4279 (**Fig. 5D-E and Fig. S7E**). These data suggest that downregulation of RAD18 may be a biologically relevant mechanism of USP1 inhibitor resistance, consistent with our RAD18 KO studies and prior reports (31, 32, 36).

Previous work has shown that USP1 inhibitors activate replication stress signaling in BRCA1-deficient cells (31, 32). We therefore reasoned that further induction of this response through an independent pathway might restore USP1 inhibitor sensitivity. Consistent with this, a sublethal dose of the ATR inhibitor M1774 (52) re-sensitized C.I4 cells to I-138 (**Fig. 5F**).

Together, these results indicate that resistance to USP1 inhibition is partially mediated by reduced RAD18-dependent PCNA ubiquitination but can be overcome by exacerbating replication stress.

## Discussion

RING finger E3 ubiquitin ligases function as substrate-specificity factors that direct their ubiquitin-charged E2 enzymes to ubiquitinate substrate lysine residues (18). Defining how E3 ligases recognize and bind their substrates remains challenging, in part due to the weak and transient nature of these interactions (53–55). RAD6-RAD18 is the principal E2-E3 pair that catalyzes mono-ubiquitination of PCNA (19). Here, we identify a PIP motif within the SAP domain of RAD18 that is critical for PCNA mono-ubiquitination. AlphaFold modelling suggests that this motif may adopt a short helical conformation between residues 5 and 7, consistent with canonical PIP motif architecture (9), despite diverging from the consensus sequence. NMR titration assays estimate that the SAP domain PIP motif binds PCNA with low affinity (Kd > 1,000 μM), substantially weaker than most characterized PIP motifs (9). Despite its low affinity, this interaction is consistent with the functional requirements of RAD18. Whereas DNA polymerases use PIP motifs to stably anchor to PCNA, the RAD18 PIP motif likely functions in transient substrate recognition, enabling dynamic regulation. This model is also consistent with the requirement for additional interactions, including binding to RPA70 and Pol η, for RAD18 recruitment to replication forks (23–25). Furthermore, because RAD18 functions as a dimer (20–22), the presence of two SAP-domain PIP motifs within the RAD6-RAD18₂ complex may increase the overall avidity of the interaction with PCNA.

Our data are consistent with prior studies. First, a C-terminal truncation mutant of RAD18 encompassing residues 16-366 was found to interact with RAD6 and PCNA, indicating that the primary PCNA-binding surface resides within this region (22). Second, the SAP domain is required for PCNA mono-ubiquitination (21, 22), and this function appears to be independent of its DNA-binding role (37). In addition, an intramolecular interaction between the SAP and RING domains was found to be essential for PCNA mono-ubiquitination (40). Third, Pol η and Pol δ, which bind the PCNA universal-binding site, disrupt RAD18-dependent PCNA mono-ubiquitination *in vitro*, suggesting that RAD18 engages the same PCNA interface through a comparatively weaker interaction (26). Identification of a PIP motif within the RAD18 SAP domain that interacts weakly with PCNA thereby provides a unifying molecular explanation for these prior experimental findings.

Having defined a required interface for RAD18-directed PCNA mono-ubiquitination, we examined its functional consequences for USP1 inhibitor sensitivity. RAD18 is a key driver of USP1 inhibitor-induced cell death in BRCA1-deficient cells (30, 32). Disruption of the RAD18-PCNA interface specifically impaired PCNA mono-ubiquitination, supporting a central role for this modification in USP1-BRCA1 synthetic lethality. Consistent with this, mutation of the SAP-domain PIP motif suppressed both USP1 inhibitor-induced accumulation of ubiquitinated PCNA and loss of fork-associated PCNA. These observations align with studies showing that mono-ubiquitinated PCNA can be extended with proteasome-targeting poly-ubiquitin chains that are normally removed by USP1 (29, 31, 36), and that loss of fork-associated PCNA contributes to ssGAP accumulation and USP1 inhibitor sensitivity (31).

Reduced fork-associated PCNA, and the resulting DNA under-replication, has been linked to synthetic lethality between USP1 and Okazaki fragment-processing factors such as FEN1 and LIG1 (36). Consistent with this, USP1 inhibition induces ssGAP accumulation and increased S-phase PARylation (32), indicating a role in maintaining replication integrity. USP1 also regulates translesion synthesis at post-replicative ssGAPs generated by PrimPol-dependent repriming downstream of DNA lesions (56, 57). Although we cannot distinguish between these sources of ssGAPs, both likely contribute to the observed phenotype. Notably, ssGAPs accumulate rapidly following USP1 inhibition, and this accumulation requires an intact RAD18 SAP-domain PIP motif. Together, these observations support a model in which, regardless of their origin, ssGAPs arise from aberrant RAD18-dependent PCNA ubiquitination and subsequent loss of fork-associated PCNA (**Fig. S8**).

Beyond genetic RAD18 loss, we demonstrate that BRCA1-deficient cells with reduced RAD18 protein levels are selected during adaptation to USP1 inhibition. Although the mechanism underlying this reduction remains unclear, stable expression of WT, but not 3A, RAD18 partially restored USP1 inhibitor sensitivity, supporting a role for RAD18 downregulation in resistance. The incomplete restoration, however, indicates that additional cellular alterations also contribute to acquired USP1 inhibitor resistance. Notably, this resistance can be overcome by inducing replication stress, for example, with an ATR inhibitor.

Together, these findings define a RAD18-PCNA interface required for PCNA mono-ubiquitination that underlies USP1 inhibitor-induced killing of BRCA1-deficient cells. This mechanistic insight may inform biomarker development and therapeutic targeting of USP1 in BRCA1-deficient tumors.

## Experimental procedures

### Mammalian cell culture and lentiviral transduction

OVCAR8, UWB1.289 and RPE-1 cells were retrieved from lab stocks, authenticated by STR profiling (ATCC # 135-XV), and confirmed to be free of mycoplasma contamination by PCR (ATCC # 30-1012K). 293T cells were recently purchased from ATCC (# CRL-3216). OVCAR8 and 293T cells were cultured in high glucose DMEM (Thermo Fisher Scientific #11965092), UWB1.289 cells were grown in a 1:1 solution of RPMI 1640 (Thermo Fisher Scientific #11875093) and MEGM (Lonza # CC-3150), and RPE-1 cells were culture in DMEM/F12, GlutaMAX (Thermo Fisher Scientific # 10565018). All media was supplemented with 5% (for UWB1.289 cells) or 10% (for OVCAR8, RPE-1 and 293T cells) fetal bovine serum (FBS, Gibco, Thermo Fisher Scientific # A5670701), and 1% penicillin-streptomycin (10,000 units mL^-1^ penicillin, 10LJmg mL^-1^ streptomycin; Gibco, Thermo Fisher Scientific # 15140122). Cells were cultured in a controlled environment that was maintained at 37 °C with a humidified atmosphere containing 5% CO_2_.

True Guide Synthetic gRNAs targeting RAD18 or USP1 were purchased from ThermoFisher Scientific, mixed with TrueCut Cas9 Protein v2 (ThermoFisher Scientific # A36498) and transfected into cells with Lipofectamine CRISPRMAX Cas9 transfection Reagent (TermoFisher Scientific #CMAX00001) as per the manufacturer’s instructions. RAD18 (exon 2) was targeted with gRNA sequence 5′-AUAGAUGAUUUGCUGCGGUG-3′ and USP1 (exon 1) was targeted with gRNA sequence 5′-CUUUCACUAGGUAUGACACC-3′. Following transfection, single OVCAR8 or RPE-1 cells were isolated by limiting dilution and expanded as monoclonal cultures. For UWB1.289 cells, monoclonal populations were isolated by plating 2,500 transfected cells on 10 cm^2^ dishes, allowing colonies to grow, and then using cloning cylinders (Corning #31668) to specifically collect and expand individual colonies. Clones containing premature stop codons that disrupt USP1 or RAD18 were initially detected by western blotting. The presence of disruptive mutations was then confirmed by PCR amplification and Sanger sequencing of the target exon. Exon 2 of RAD18 was amplified using the primers 5′-CACCCAATAGTGACTATAGGCTAGG-3′ and 5′-AAGTACTACTTGGTGGAACCACC-3′, and sequenced using primer 5′-CCTTGCAGTTTATCTGGAGTTAGC-3′. Exon 1 of USP1 was amplified with primers 5′-GAGTGCTTATTGGCAGGC-3′ and 5′-CTTGAGAATCTGTGAAATCCAAAGC-3′, and sequenced using the former.

Lentiviral vectors were transduced using an established protocol (58). Briefly, lentiviral expression, packaging, and envelope vectors were transfected into 293T cells using a CalPhos mammalian transfection kit (Takara # 631312). 293T cells were grown for 48 hours, then the lentivirus-containing media collected and added to cultures of the target OVCAR8, UWB1.289 or RPE-1 cell lines. The media of target cells were also treated with 8 μg ml^-2^ polybrene infection/transfection reagent (MilliporeSigma # TR-1003-G). Transduced cells were grown for 48-72 hours, then those stably expressing the lentiviral expression vectors selected for with puromycin (2 μg ml^-2^ for OVCAR8 and UWB1.289, 20 μg ml^-2^ for RPE-1).

### Immunoblotting

Cell lysis and immunoblotting was performed as described previously (59). In brief, 20 μg of whole cell lysate was typically electrophoresed on 15-well 1.5 mm 4–12% Bis-Tris NuPage precast gels (Thermo Fisher Scientific) and transferred to nitrocellulose membranes. Membranes were blocked in Intercept (TBS) Blocking Buffer (Li-Cor # 927-60001) for 45 min, prior to incubation with primary antibodies overnight at 4 °C. The following primary antibodies were purchased from Cell Signaling Technology: Rb-α-RAD18 (clone D2B8; # 9040), Rb-α-USP1 (clone D37B4; # 8033), Rb-α-β-Actin (clone 13E5; # 4970), Ms-α-PCNA (clone PC10; # 2586), Rb-α-ubiquityl-PCNA (Lys164) (clone D5C7P; # 13439), Ms-α-Chk1 (clone 2G1D5; # 2360), Rb-α-pS345 Chk1 (clone 133D3; # 2348). Antibodies against KAP1 (Ms clone 20C1; # ab22553) and pS824 KAP1 (Rb pAb; # ab10484) were purchased from Abcam. Primary antibodies were detected using IRDye 680RD or 800CW-conjugated donkey anti-mouse or anti-rabbit fluorescent secondary antibodies (Li-Cor) and visualized using an Odyssey CLX infrared imaging system (Li-Cor). Immunoblots were quantified using Image Studio software (Li-Cor).

### Protein expression and purification

*Escherichia coli* BL21(DE3) cells transformed with the PCNA plasmid were grown at 37°C in 100% D_2_O-based M9 minimal medium supplemented with ^15^NH_4_Cl as a sole source of nitrogen, until reaching the OD_600_ of ∼1.0. Expression of the ^15^N/^2^H-labeled protein was induced by adding 1 mM Isopropyl β-D-1-thiogalactopyranoside (IPTG) overnight at 20°C.

Cells were harvested by centrifugation, resuspended in a buffer containing 20 mM phosphate buffer pH 8, 250 mM NaCl, 10 mM imidazole, 1 mM PMSF, lysed by sonication, and centrifuged at 15,000 rpm for 1 hour. The supernatant was filtered and applied to a TALON HisPur cobalt resin (Thermo Scientific). The protein was eluted in a buffer containing 20 mM phosphate pH 8, 250 mM NaCl and 300 mM imidazole. Thrombin protease was then added to the sample overnight at 4°C to remove the 6-His tag. The protein sample was further purified by size-exclusion chromatography on a HiLoad Superdex 200 column (Cytiva). NMR buffer contained 50 mM Na_2_HPO_4_ pH 7.0, 100 mM NaCl, 2 mM dithiothreitol (DTT) and 10% (v/v) D_2_O. Final NMR samples of PCNA contained ∼0.4 mM protein. Prior to NMR experiments, protein samples were incubated at room temperature for at least 4 days to allow for ^2^H/^1^H exchange of the protein amide groups.

Peptides representing RAD18 residues 235-265 (SSVHKRKPLPKTVYNLLSDRDLKKKLKEHGL) and 241-254 (WT = KPLPKTVYNLLSDR; V247A = KPLPKT<u>A</u>YNLLSDR; L250A_L251A = KPLPKTVYN<u>AA</u>SDR; V247A_L250A_L251A = KPLPKT<u>A</u>YN<u>AA</u>SDR) were chemically synthesized by GenScript at a purity of ≥90%. The lyophilized peptides were resuspended in a buffer containing 50 mM Na_2_HPO_4_ pH 7.0, 100 mM NaCl and 2 mM DTT to create 10 mM stock solutions.

### NMR titration experiments

All NMR experiments were collected on an 800 MHz (^1^H) Bruker Avance Neo spectrometer, equipped with a cryogenic probe, at 35°C. ^15^N/^2^H-labeled PCNA (427 μM) was titrated with unlabeled WT Rad18 peptide (235–265) up to a 6-fold peptide excess. Separately, unlabeled Rad18 peptides (241-254 (WT); 241-254 (V247A); 241-254 (L250A_L251A); or 241-254 (V247A_L250A_L251A)) were titrated into ^15^N/^2^H-labeled PCNA (355 μM) to an 8-fold peptide excess. ^1^H-^15^N TRSOY spectra were collected at each titration point to monitor binding.

Data were processed using NMRPipe (60) and analysed using Sparky (61). All data processing and analysis were performed on the NMRbox (62) platform. Previously reported NMR resonance assignment of PCNA (BMRB 15501) (43) was used to assign the peaks in the PCNA spectra. Per-residue NMR chemical shift perturbations (Δω_obs_) were calculated as Δω_obs_ = (Δω_N_^2^ + Δω_H_^2^)^1/2^ where Δω_N_ and Δω_H_ are the chemical shift differences between free and bound samples measured in Hz for ^15^N and ^1^H, respectively. The resulting Δω_obs_ values measured in ^1^H ppm were mapped onto the PCNA structure (PDB: 4RJF) (44) to visualize the binding interfaces.

NMR titrations of ^15^N/^2^H PCNA with unlabeled RAD18 peptides were used to estimate and compare their binding affinities (K_D_) using the following equation:

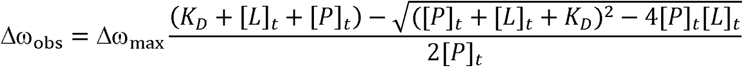

Where [P]_t_ and [L]_t_ are the total protein and ligand concentrations and Δω_max_ is the chemical shift difference between free and RAD18-bound PCNA at saturation. A representative peak (I128) in the PCNA ^15^N TROSY spectrum was used. The titration curves were fit assuming a shared Δω_max_ using Prism V10.4.1 (GraphPad).

### Protein structure prediction and modelling

AlphaFold3 and the AlphaFold Server (63) were used to model the predicted interaction between the RAD18 SAP domain PIP motif and PCNA. Models were visualized in PyMOL V3.0.4 (Schrödinger) using the top-ranking predictions per seed. An AlphaFold model of full-length RAD18 (AF-Q9NS91-F1-v4) was retrieved from the AlphaFold Protein Structure Database (42).

### Inhibitors and DNA damage

The USP1 inhibitors, I-138 (# E1479) and KSQ-4279 (# E1214), as well as the ATR inhibitor, M1774 (tuvusertib; # E1411), were purchased from SelleckChem. Inhibitors were dissolved in DMSO at concentrations of 10 μM (for KSQ-4279 and M1774) or 20 μM (for I-138) and stored in aliquots at −80 °C.

UV-C irradiation of cells was performed with a portable UVP 3UV lamp (Analytik Jena # 95-0343-02) that was suspended ∼900 mm from the irradiation surface. UV-C was emitted at 250 nm and irradiated the surface at an intensity of 40 μW / cm^2^. Cells were irradiated with 20 J/m^2^ of UV-C (50 second exposure) to induce PCNA mono-ubiquitination. Cell culture media was removed immediately prior to irradiation and replaced afterwards.

### Clonogenic Assays

UWB1.289-derived cell lines were seeded into 6-well plates at densities of 2,500 cells per well. Cells were allowed to settle for ∼4 hours and then treated with I-138, KSQ-4279, or M1774. For M1774 and I-138 co-treatment, cells were resuspended and plated in 0.5 nM of M1774, allowed to settle, and then treated with I-138. Cells were then grown for 10-14 days to allow colonies to form. Media was subsequently removed, and colonies were fixed in methanol for 1 hour, stained with crystal violet (2.5% (w/v) crystal violet, 25% ethanol (v/v)) for 1 h, and plates were washed and dried. Plates were then scanned using an Oxford Optronix GelCount imaging platform, and clonogenic growth quantified using GelCount software (version 1.4.1.11). Clonogenic growth is reported as Total Colony Volume, calculated as the sum of per colony volume values (the product of optical density x area values). Three technical replicates of each cell line were included per dose. The averages of three biological replicates were graphed using Prism 10 (GraphPad).

### DNA Fiber Assay with S1 Nuclease Digestion

DNA fiber assays to quantify ssDNA gaps were performed as previously described (32, 64). Briefly, cells were seeded 24 h prior to labeling. Cells were first labeled with 200 μM CldU (Sigma, cat # C6891) for 30 min, washed three times with PBS, and then labeled with 100 μM IdU (Sigma, cat # I7125) for 2 h. DMSO or 0.5 μM I-138 were added concurrently with IdU. After labeling, cells were washed with PBS and permeabilized in CSK buffer (100 mM NaCl, 10 mM MOPS, 3 mM MgCl2, 300 mM sucrose, 0.5% Triton X-100) for 10 min at room temperature. Nuclei were washed once with PBS, incubated briefly in S1 buffer (50 mM NaCl, 300 mM sodium acetate, pH 4.6, 10 mM zinc acetate, 5% glycerol), and then treated with 10 U/ml S1 nuclease (Sigma, cat # N5661) in S1 buffer for 30 min at 37°C. Nuclei were collected in 1 ml PBS (0.1% BSA), centrifuged (5 min, 1500 rpm), and resuspended in PBS. DNA was isolated via agarose plug preparation and proteinase K treatment overnight at 50°C. Agarose plugs were digested with β-agarose overnight, and DNA was combed onto silanized coverslips using the Molecular Combing System (Genomic Vision). DNA was denatured in 0.5 M NaOH/1 M NaCl for 8 min and dehydrated through an ethanol series (70%, 90%, 100%, 3 min each). Fibers were immunostained using rat-α-BrdU (CldU-specific; Abcam cat # ab6326), mouse-α-BrdU (IdU-specific; BD Biosciences, cat # 347580), and mouse-α-anti-ssDNA (Developmental Studies Hybridoma Bank, University of Iowa). Secondary antibodies included Cy5® goat-α-rat (Abcam cat# ab6565), Cy3® goat-α-mouse (Abcam, cat# ab97035), and BV480 goat-α-mouse (BD Biosciences, cat# 564877). Fibers were scanned using the FiberVision S system and analyzed in ImageJ. IdU track lengths were quantified, and results graphed using Prism software 10 (GraphPad).

### SIRF assay

SIRF assay was performed as previously described (32, 64). Cells were seeded on coverslips and incubated overnight. The following day, cells were treated with 0.5 μM I-138 for the indicated times, and EdU was added for the final 15 min of incubation. Coverslips were washed with PBS, and cells were pre-extracted with 0.5% Triton X-100 for 2 min. Cells were then fixed with 4% paraformaldehyde (PFA) for 10 min at room temperature and washed three times with PBS. Permeabilization was performed with 0.2% NP-40 in PBS for 2 min, followed by blocking in 3% BSA in 0.1% Triton X-100. Cells were incubated with freshly prepared click reaction mix and incubated for 30 min. Coverslips were washed three times with PBS and incubated overnight at 4 °C with anti-ub-PCNA or anti-PCNA with anti-biotin antibodies. Coverslips were then washed in Wash Buffer A for 5 min each. PLA was performed using the Duolink® In Situ Detection Kit (Sigma-Aldrich # DUO92008) according to the manufacturer’s instructions. Briefly, Duolink In Situ PLA probes (anti-mouse PLUS and anti-rabbit MINUS) were diluted 1:5 in blocking buffer and applied to coverslips (25 μl/well) for 1 h at 37 °C. After washing in Wash Buffer A, ligation was carried out using Duolink ligation stock (1:5) and ligase (1:40) in high-purity water for 30 min at 37 °C. Coverslips were washed twice for 2 min each in Wash Buffer A. Amplification was carried out using amplification stock (1:5) and polymerase (1:80) in high-purity water) for 1 h 40 min at 37 °C. After amplification, coverslips were washed twice for 10 min each in Wash Buffer B, followed by one wash in 0.01× diluted Wash Buffer B. Coverslips were mounted with ProLong™ Diamond Antifade Mountant with DAPI (Thermo Fisher). Images were acquired using Zeiss AxioObserver 7 inverted widefield microscope equipped with ApoTome.2, X-Cite 120 Boost LED System, Axiocam 506 mono camera and ZEN image acquisition software. Objective 63X oil: Plan-Apochromat 63x/1.40 Oil WD=0.19mm M27; or Objective 20X air: Plan-Apochromat20x/0.8 M27 FWD=0.55mm were used. Images were analyzed using CellProfiler software. To control for differences in EdU incorporation across samples, protein-biotin foci counts were normalized to the corresponding average biotin-biotin foci within each sample.

## Data Availability

All study data are included in the article and/or *supporting information*. Uncropped immunoblot images, clonogenic and PLA assay quantification and generated AlphaFold models are available from Mendeley Data (https://data.mendeley.com/datasets/wv5p74n2vj). Further information and requests for resources and reagents should be directed to ADA.

## Supporting Information

This article contains supporting information.

## Acknowledgments

We would like to thank all members of the D’Andrea laboratory for their helpful suggestions and comments. We acknowledge use of the web portals from the FP7 WeNMR, H2020 West-Life, EOSC-hub, and EGI-ACE European e-Infrastructure projects, and recognize their supporting organizations and national GRID Initiatives, for their vital contributions to the EGI infrastructure.

## Author Contributions

**Nicholas W. Ashton**: Conceptualization, Methodology, Investigation, Formal analysis, Visualization, Writing – original draft, Writing – review & editing. **Ramya Ravindranathan**: Methodology, Investigation, Formal analysis, Writing – original draft. **Emilie J. Korchak**: Investigation, Formal analysis, Writing – original draft. **Ozge S. Somuncu**: Investigation; **Gabriella A. Zambrano**: Investigation. **Sirisha V. Mukkavalli:** Project administration. **Shuhei Asada**: Investigation. **Dmitry Korzhnev**: Formal analysis. **Irina Bezsonova**: Conceptualization, Supervision, Funding acquisition, Formal analysis, Writing – review & editing. **Alan D. D’Andrea**: Conceptualization, Supervision, Funding acquisition, Writing – review & editing.

## Funding and additional information

This work was supported by NIH grants to ADD (R01CA31133) and IB (R35GM156397), as well as by funding from the Ludwig Center at Harvard.

## Conflict of interest

A.D.D. reports consulting for Bayer AG, Bristol Myers Squibb, EMD Serono, Impact Therapeutics, Tango Therapeutics, Roche Pharma, and Covant, Therapeutics; is an Advisory Board member for Impact Therapeutics; and reports receiving commercial research grants from EMD Serono, Moderna, and Tango Therapeutics.

## Abbreviations and nomenclature

ATR: Ataxia telangiectasia and Rad3-related
BSA: bovine serum albumin
BRCA1: breast cancer type 1 susceptibility protein
CldU: chlorodeoxyuridine
EdU: 5-ethynyl-2′-deoxyuridine
FBS: fetal bovine serum
FEN1: flap endonuclease 1
HR: homologous recombination
IdU: iododeoxyuridine
IPTG: isopropyl β-D-1-thiogalactopyranoside
KO: knockout
LIG1: DNA ligase 1
PARylation: poly(ADP-ribose)ylation
PCNA: proliferating cell nuclear antigen
PFA: paraformaldehyde
PIP: PCNA-interacting peptide
PLA: proximity ligation assay
Pol η: DNA polymerase eta
Pol ι: DNA polymerase iota
Pol δ: DNA polymerase delta
Pol ε: DNA polymerase epsilon
PRIMPOL: primase-polymerase
RING: really interesting new gene
RPA: replication protein A
SAP: SAF-A/B, Acinus and PIAS
SEM: standard error of the mean
SIRF: in situ analysis of protein interactions at DNA replication forks
ssDNA: single-stranded DNA
ssGAP: single-stranded DNA gap
TLS: translesion synthesis
UAF1: USP1-associated factor 1
UBZ: ubiquitin-binding zinc finger
ub-PCNA: mono-ubiquitinated PCNA
USP1: ubiquitin-specific protease 1
UV-C: ultraviolet C

## Notes

### Summary of Updates

We have altered the title of the manuscript, replaced figures 3E and 5C, and added figure 3F. We have also refined various sections of the text.

