## Supporting information for "USP1 inhibition promotes RAD18-dependent PCNA degradation and BRCA1 synthetic lethality"

**Running title:** RAD18 and USP1-BRCA1 Synthetic Lethality

**Keywords:** ubiquitin-specific protease 1; E3 ubiquitin-protein ligase RAD18; PCNA-interacting peptide; BRCA1-deficient cancers; synthetic lethality

SI Figures

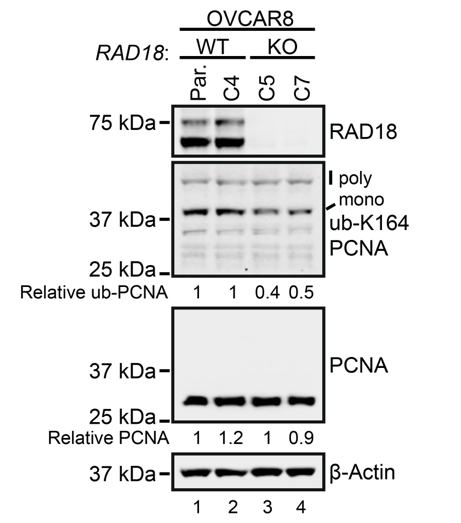

Figure S1. Loss of RAD18 reduces basal PCNA mono-ubiquitination without altering total PCNA levels in OVCAR8 cells. Immunoblot of whole-cell lysates from parental OVCAR8 cells (Par.), a WT RAD18 clone (C4), and two RAD18 knockout clones (C5 and C7). Total PCNA levels were normalized to β-Actin. Mono-ubiquitinated PCNA (ub-PCNA) levels were normalized to total PCNA. Values are expressed relative to parental OVCAR8 cells (lane 1).

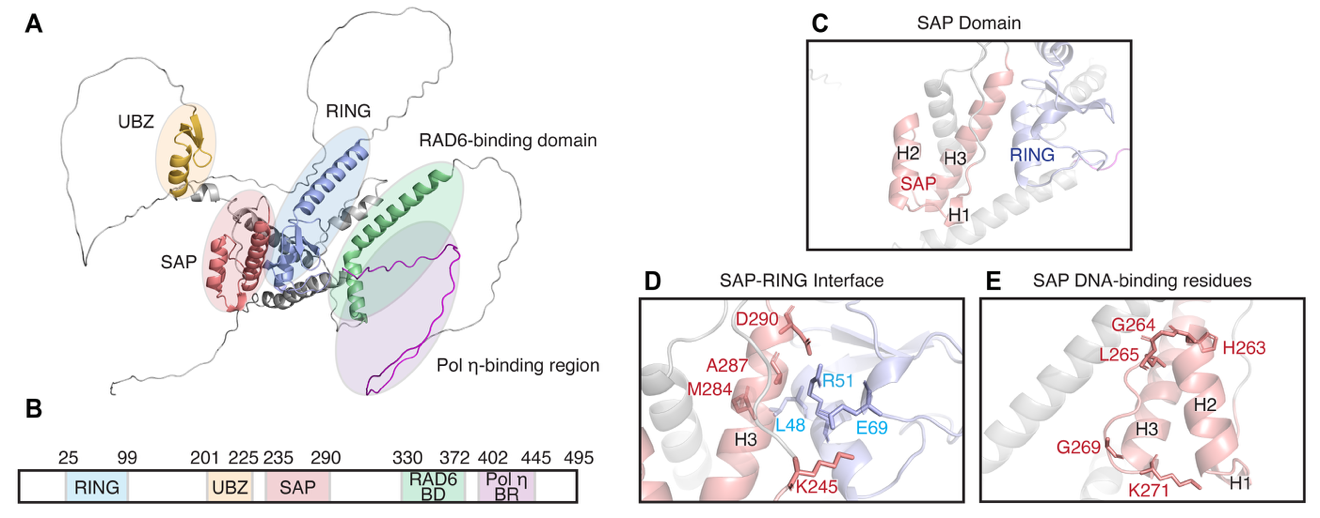

Figure S2. AlphaFold structural prediction of RAD18 domain organization. (A) AlphaFold model of RAD18 retrieved from the AlphaFold Protein Structure Database (1) (AF-Q9NS91-F1-v4). Colors indicate the RING, UBZ, SAP and RAD6-binding domains, as well as the Pol η-binding region. (B) RAD18 domain architecture schematic (reproduced from Figure 2A). (C) Enlarged view of the SAP domain, indicating the three helices (H1, H2, H3), as well as the proximity of the SAP and RING domains. (D) Enlarged view of the interface between H3 of the SAP domain and the RING finger domain. The residues highlighted are those identified as important binding residues (2). (E) Enlarged view of the SAP domain, highlighting residues implicated in DNA-binding within the loop between H2 and H3 (3, 4).

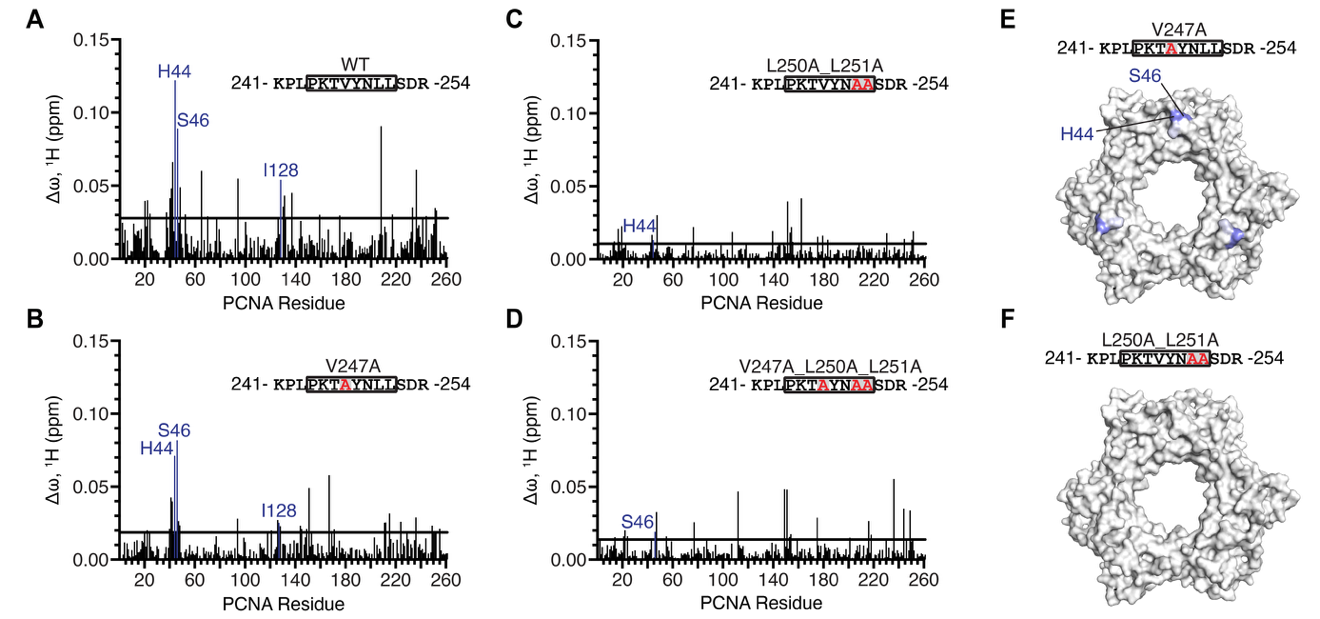

Figure S3. The RAD18 SAP domain contains a PIP motif that binds the PCNA universal binding site. (A-D) Bar graphs showing per-residue chemical shift perturbations (Δω) in the ^15^N-TROSY spectrum of ^15^N/^2^H-labeled PCNA following addition of unlabeled RAD18 peptides (residues 241-254). (A) = WT, (B) = V247A, (C) = L250A_L251A, (D) = V247A_L250A_L251A. PCNA universal binding site residues are highlighted in blue and labeled. (E-F) Mapping of PCNA-binding sites for RAD18 mutant peptides V247A (E) and L250A_L251A (F) onto the surface of PCNA (PDB: 4RJF) (5). Surfaces are color-coded according to Δω values from smallest (white) to largest (blue). Universal binding site residues are highlighted and labeled in (E). Mapping for the WT and V247A_ L250A_L251A peptides are shown as Figure 2D.

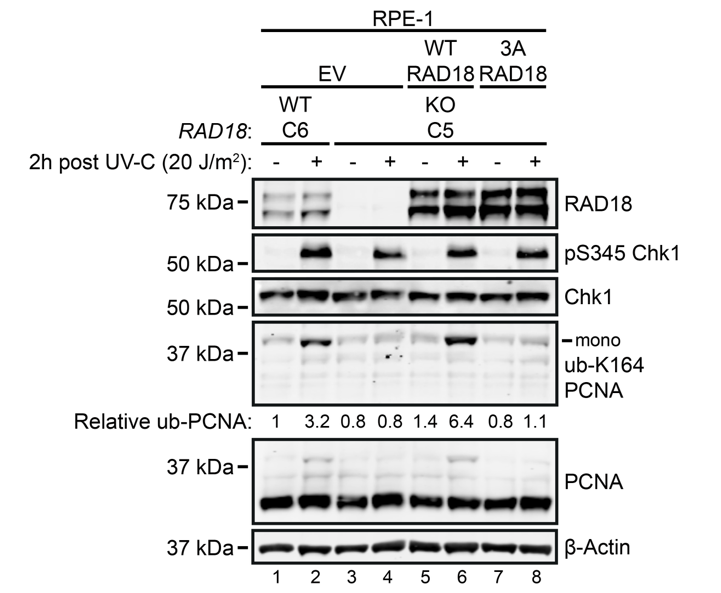

Figure S4. Mutation of the RAD18 SAP domain PIP motif disrupts UV-C-induced PCNA mono-ubiquitination in RPE-1 cells. Immunoblot analysis of UV-C-induced PCNA mono-ubiquitination in WT RAD18 RPE-1 cells (C4) and RAD18 KO RPE-1 cells (C5) stably expressing WT RAD18, 3A RAD18 (V247A_L250A_L251A), or empty vector (EV). Cells were exposed to 20 J m⁻² UV-C or left untreated and harvested after 2 h. Relative mono-ubiquitinated PCNA (ub-PCNA) levels were calculated as the ub-PCNA:PCNA ratio and expressed relative to the untreated WT RAD18 clone (lane 1).

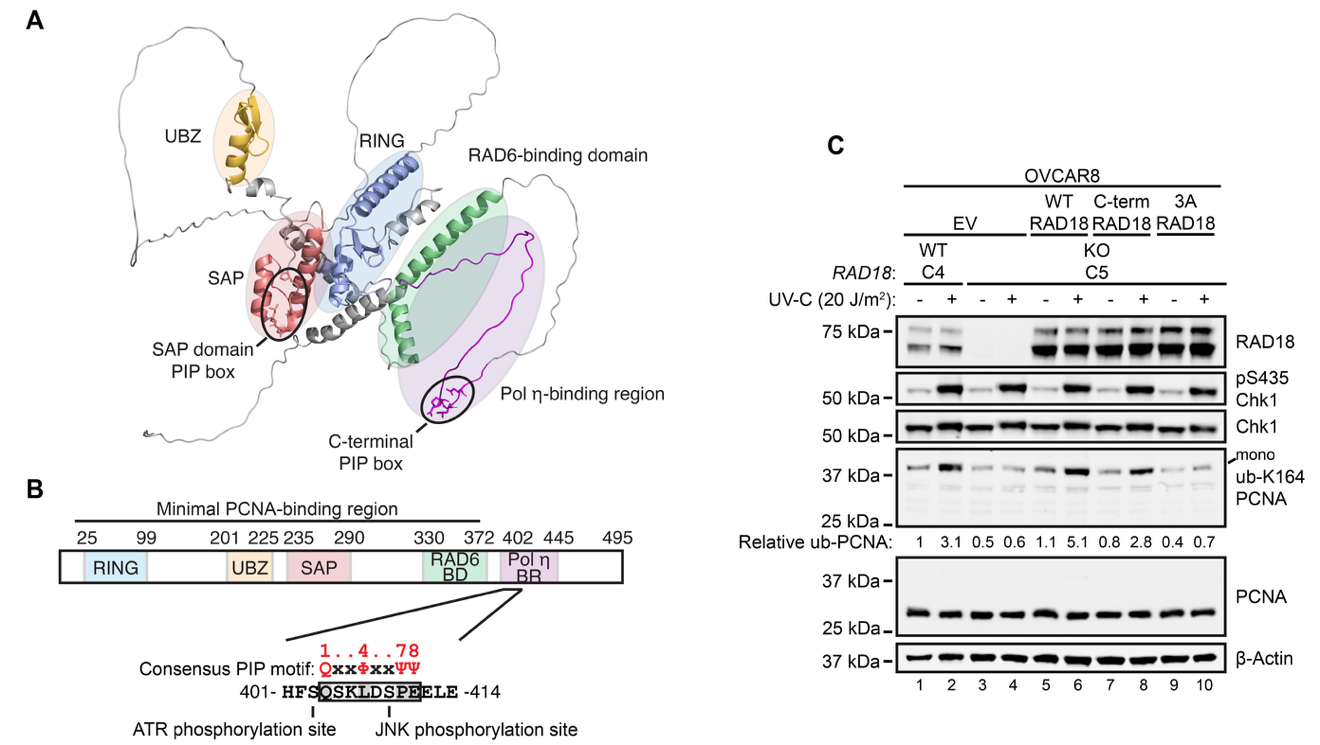

**Figure S5. A C-terminal RAD18 motif contributes to UV-C-induced PCNA mono-ubiquitination.** (**A**) AlphaFold model of RAD18 (also appears as Figure S2A), highlighting the SAP domain PIP motif and a putative C-terminal PIP motif **(6)**. (**B**) Schematic of RAD18 domain architecture (also appears as Figure 2A), indicating the putative C-terminal PIP motif and adjacent ATR and JNK phosphorylation sites. (**C**) Immunoblot analysis of UV-C-induced PCNA mono-ubiquitination in WT RAD18 OVCAR8 cells (C4) and a RAD18 KO OVCAR8 clone (C5) stably expressing WT RAD18, 3A RAD18 (V247A_L250A_L251A), the C-terminal PIP mutant (Q404A_L407A), or an empty vector (EV). Cells were exposed to 20 J m^-2^ UV-C or left untreated and harvested after 2 hours. Relative ub-PCNA levels were calculated based the ub-PCNA:PCNA ratio and expressed relative to the untreated WT RAD18 clone (lane 1).

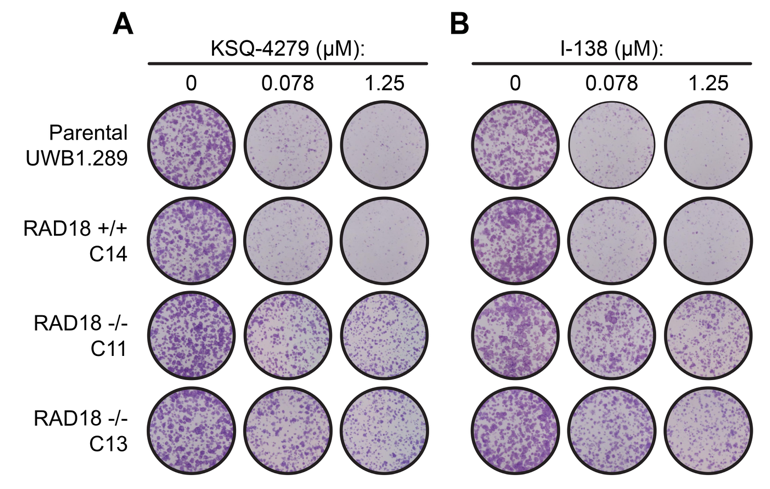

Figure S6. RAD18 KO UWB1.289 cells exhibit reduced sensitivity to USP1 inhibitors. (A-B) Representative images of clonogenic survival of parental UWB1.289 cells, a WT RAD18 clone (C14), and two RAD18 knockout clones (C11 and C13), treated with the USP1 inhibitor, I-138 (A) or KSQ-4279 (B). WT RAD18 cells were stained after 10 days of treatment, while RAD18 KO clones were stained after 14 days. Values derived from these images are graphed in Fig. 4B and C.

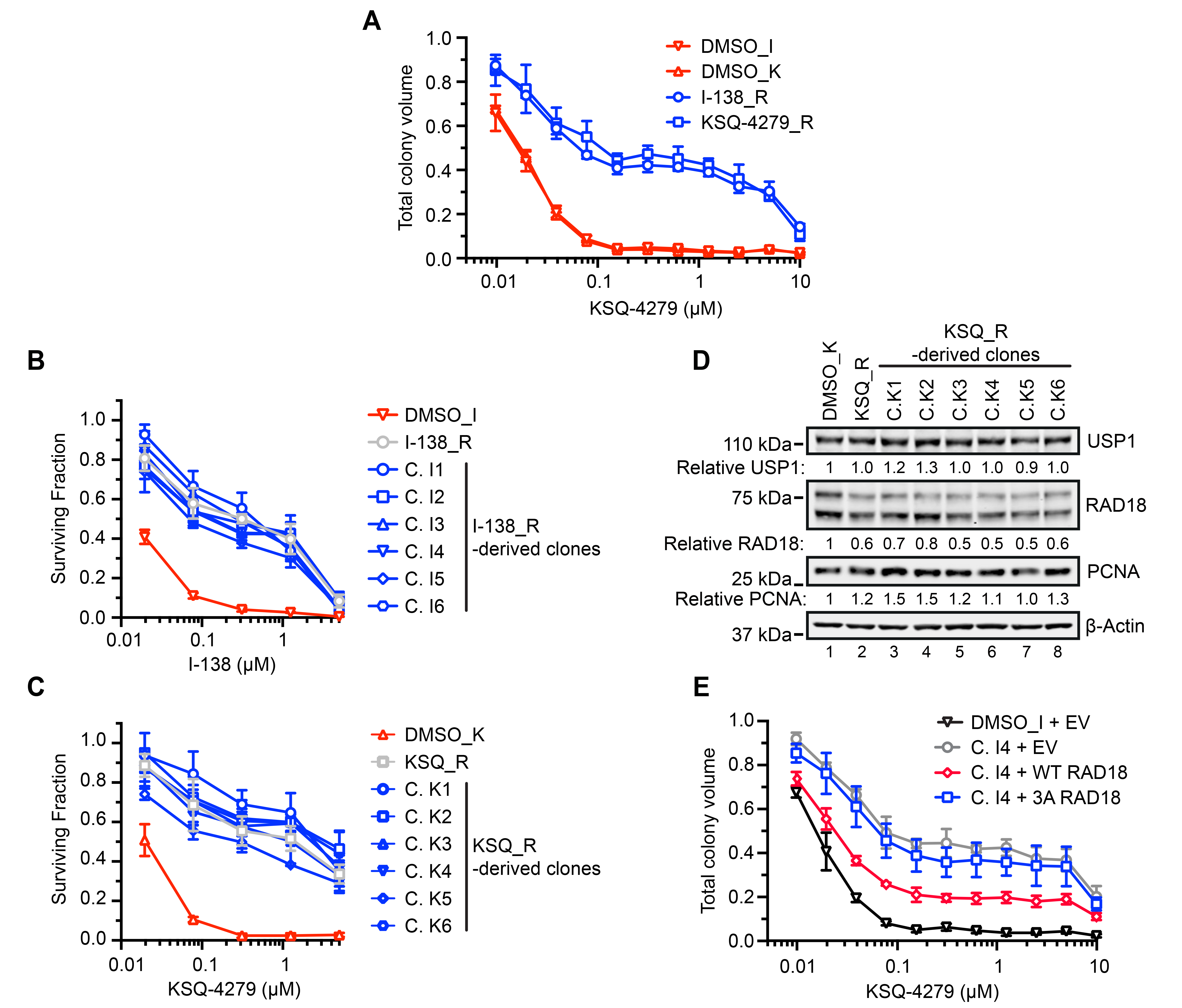

Figure S7. Clones derived USP1 inhibitor-adapted UWB1.289 populations exhibit comparable resistance. (A) Clonogenic survival of the populations in Fig. 5A following treatment with KSQ-4279. Adapted populations were maintained drug-free for ≥1 week before analysis. Values represent means of three biological replicates ± SD. (B) Clonogenic survival following I-138 treatment in control UWB1.289 cells (DMSO_I), a population adapted by culture for ~4 months in progressively increasing I-138 concentrations (I-138_R), and six clones derived from the I-138_R population. (C) Clonogenic survival following KSQ-4279 treatment in control UWB1.289 cells (DMSO_K), a population adapted by culture for ~4 months in progressively increasing KSQ-4279 concentrations (KSQ_R), and six clones derived from the KSQ_R population. For (B) and (C), values represent means of three biological replicates, with clonogenic survival calculated relative to DMSO-treated cells. Error bars indicate SD. (D) Immunoblot of whole-cell lysates from DMSO_K, KSQ_R, and derived clones K1-I6. USP1, RAD18, and PCNA levels were normalized to β-Actin and expressed relative to lane 1. (E) Clonogenic survival of cell lines in Fig. 5D following treatment with KSQ-4279. Values represent means of three biological replicates normalized to DMSO controls ± SD.

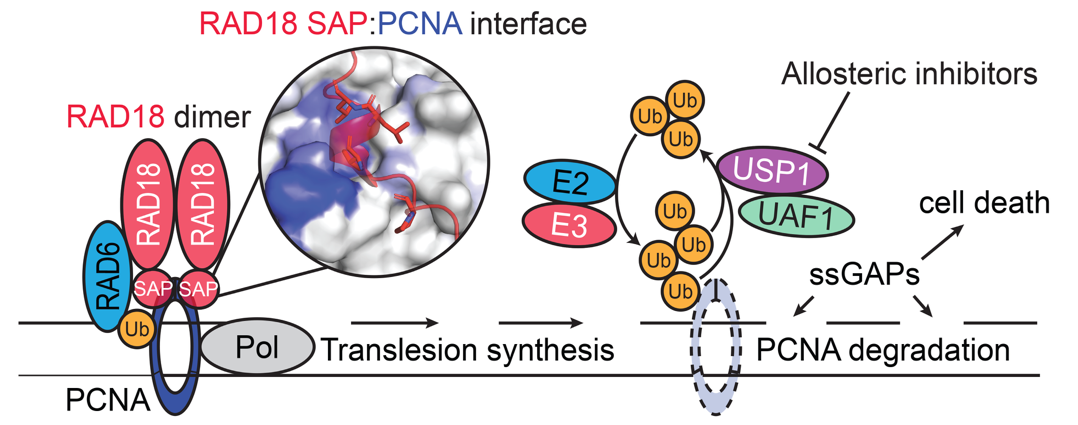

Figure S8. A RAD18-PCNA interface enables PCNA mono-ubiquitination and USP1-BRCA1 synthetic lethality. A model depicting the contribution of PCNA mono-ubiquitination to USP1 inhibitor-induced PCNA degradation, ssGAP accumulation and cell death. The inset image of the RAD18 SAP PIP motif binding to the universal binding site of PCNA is derived from Fig. 3B.
